# beachmat: a Bioconductor C++ API for accessing single-cell genomics data from a variety of R matrix types

**DOI:** 10.1101/167445

**Authors:** Aaron T. L. Lun, Hervé Pagès, Mike L. Smith

## Abstract

Recent advances in single-cell RNA sequencing have dramatically increased the number of cells that can be profiled in a single experiment. This provides unparalleled resolution to study cellular heterogeneity within biological processes such as differentiation. However, the explosion of data that are generated from such experiments poses a challenge to the existing computational infrastructure for statistical data analysis. In particular, large matrices holding expression values for each gene in each cell require sparse or file-backed representations for manipulation with the popular R programming language. Here, we describe a C++ interface named *beachmat*, which enables agnostic data access from various matrix representations. This allows package developers to write efficient C++ code that is interoperable with simple, sparse and HDF5-backed matrices, amongst others. We perform simulations to examine the performance of *beachmat* on each matrix representation, and we demonstrate how *beachmat* can be incorporated into the code of other packages to drive analyses of a very large single-cell data set.

## Introduction

Recent advances in single-cell RNA sequencing (scRNA-seq) technologies have led to an explosion in the quantity of data that can be generated in routine experiments. Droplet-based methods such as Drop-Seq [16], inDrop [12] and GemCode [25] allow expression profiles to be captured for each of thousands to millions of cells. It hardly needs to be said that this is a substantial amount of data – the expression profile for each cell consists of a measure of expression for each transcriptionally active (and polyadenylated) genomic feature, of which there are usually 10,000-40,000 in the current genome annotations for most model organisms. Careful computational analysis is critical to extract meaningful biology from these data, but the sheer volume strains existing pipelines and methods designed for single-cell data processing. The challenge is compounded by the presence of large-scale projects such as the Human Cell Atlas [19], which aims to use single-cell ‘omics to profile every cell type in the human body. Similar issues are encountered outside of transcriptomics, with single-cell ATAC-seq [3] and bisulfite sequencing [22] providing region- to base-level resolution of biochemical events (chromatin accessibility and DNA methylation, respectively). This results in even more data compared to gene-level expression values.

It is fair to say that the R programming language [18] is the premier tool of choice for statistical data analysis. R provides well-designed, rigorously-tested implementations of a large variety of statistical methods. Its interactive nature makes it easy for newcomers to learn and lends itself to data exploration and research, while its programming features allow more experienced users to readily assemble complex analyses. It is also extensible through the installation of optional packages, often contributed by the research community, which provide implementations of bespoke methods targeted to specific scientific problems. In particular, the Bioconductor project [6] supports a number of packages for biological data analysis, many of which focus on the processing of genomics data [9]. Packages are usually written in R but can also include compiled code (e.g., in C/C++ or Fortran), which is beneficial for computationally intensive tasks where high performance is required. For C++ code, this process is facilitated by the *Rcpp* package [4], which simplifies the integration of package code with the R application programming interface (API).

In its simplest form, a scRNA-seq data set consists of a count matrix where each column is a cell, each row is a gene, and the value of each matrix entry is set to the quantified expression (e.g., number of mapped reads, transcripts-per-million) for that gene in that cell. This can be most directly represented in R as a simple matrix, where each entry is explicitly stored in memory. Alternatively, it can be represented as a sparse matrix using classes from the *Matrix* package [1], which saves memory by only storing non-zero entries. This exploits the fact that scRNA-seq protocols have low capture efficiencies [7] – RNA molecules are present in cells but are not reverse-transcribed to cDNA for sequencing, resulting in a preponderance of zeroes in the final count matrix. Droplet-based protocols also exhibit very low sequencing depth per cell, further increasing the sparsity of the data. Another option is to use file-backed representations such as those in the *bigmemory* [10] or *HDF5Array* packages, where the data set is stored on disk and parts of it are extracted into memory upon request. In each case, methods are provided in R for common operations such as subsetting, transposition and arithmetic, such that code written by users (or other developers) can be agnostic to the exact representation of the matrix. This simplifies the development process and improves interoperability.

Unfortunately, for compiled code written in statically typed languages like C++, the details of the matrix representation must be known during compilation. This makes it difficult to write a single, general piece of code that can be applied to many different representations. Writing multiple versions for each representation is difficult and unsustainable when more representations become available. The alternative is to perform all processing in R to exploit the availability of common methods. However, this is an unappealing option for high-performance code. For scRNA-seq data stored in matrices, consider the most common access pattern, i.e., looping across all cells or genes and performing operations on the cell- or gene-specific expression profiles. If this was performed in R, the code within the loop would need to be re-interpreted at each of thousands or millions of iterations. This increases the computational time required to perform analyses, which is inconvenient for small scripts, undesirable for interactive analyses and unacceptable for large simulation studies. It would clearly be preferable to implement critical functions (including loops) in compiled code wherever possible.

Here, we describe a C++ API named *beachmat* (using Bioconductor to handle Each Matrix Type), which enables access to R matrix data in a manner that is agnostic to the exact matrix representation. This allows developers to implement computationally intensive algorithms in C++ that can be immediately applied to a wide range of R matrix classes, including simple matrices, sparse matrices from the *Matrix* package, and HDF5-backed matrices from the *HDF5Array* package. Using simulated and real scRNA-seq data, we assess the performance of *beachmat* for data access from each matrix representation. We show that each representation has specific strengths and weaknesses, with a clear memory-speed trade-off that motivates the use of different representations in different settings. We also demonstrate how *beachmat* can be used by other Bioconductor packages to empower the analysis of a very large scRNA-seq data set. By operating synergistically with existing Bioconductor infrastructure, *beachmat* extends R’s capabilities for analyzing scRNA-seq and other large matrix data.

## Description of the *beachmat* API

### Overview of the API

The *beachmat* API uses C++ classes to provide a common interface for data access from R matrix representations. We define a base class that implements common methods for all matrix representations. Each specific representation is associated with a derived C++ class that provides customized implementations of the access methods. The intention is for a user to pass in an R matrix of any type, in the form of an **RObject** instance from the *Rcpp* API (Figure 1). A function is then called to produce its C++ equivalent, returning a pointer to the base class. This pointer is the same regardless of the R representation and can be used in downstream code to achieve run-time polymorphism.

**Figure 1.**
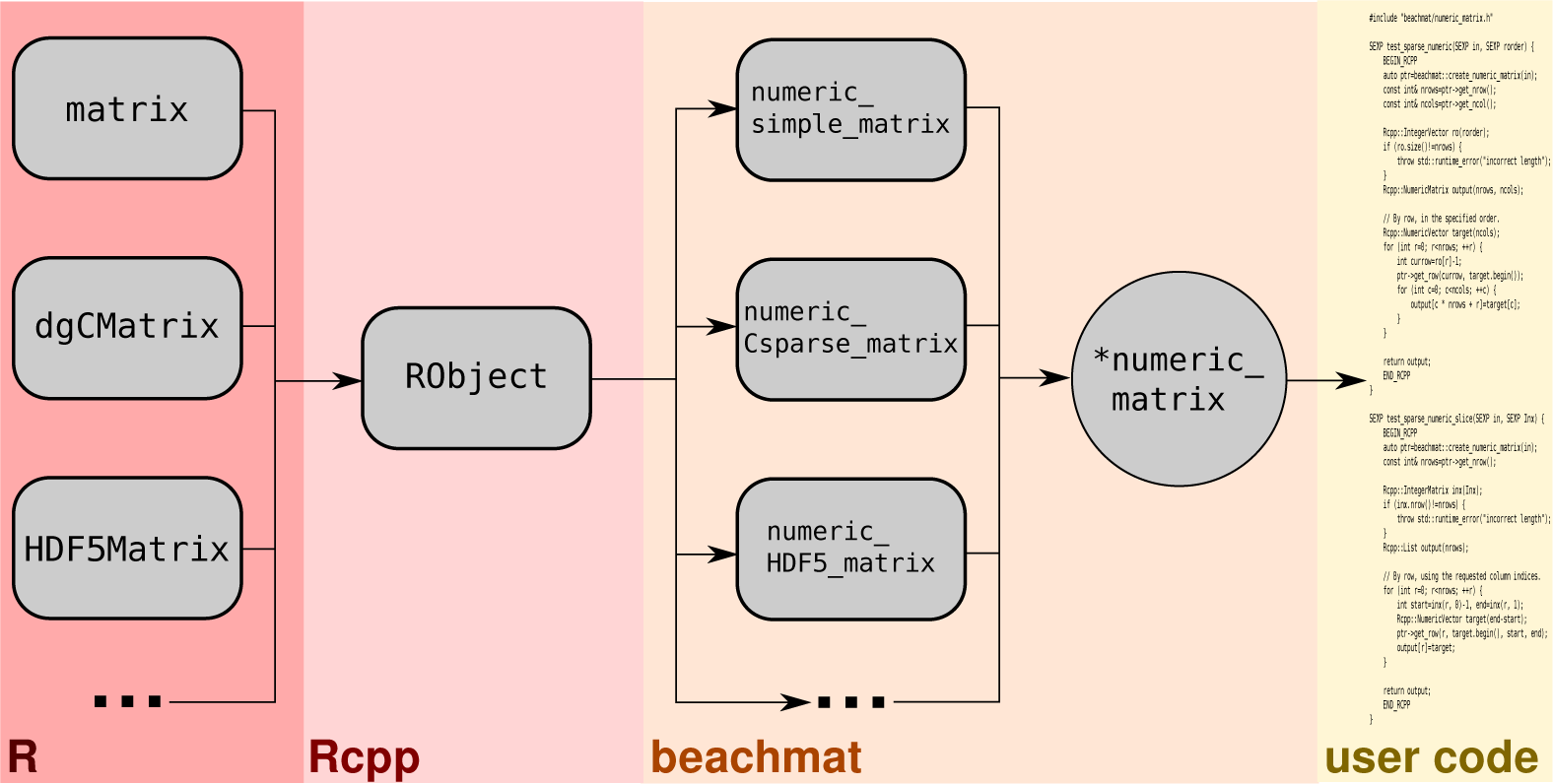
Schematic of the *beachmat* workflow. Various matrix representations at the R level are passed as **RObject** instances to a C++ function. *beachmat* identifies the specific representation, constructs an instance of the appropriate C++ derived class, and returns a pointer to base class. (In this case, a **numeric_matrix** pointer is returned for input matrices holding double-precision data.) This pointer can then be used in user-level code in a manner that is agnostic to the details of the original representation.

While the API is agnostic to the matrix representation, it still needs to know the type of data that are stored in the matrix. We use C++ templating to recycle the code to define specific classes for common data types, i.e., logical, integer, double-precision floating point or character strings. The same methods are available for all classes of each data type, improving their ease of use for developers. Briefly, when access to a specific row or column (or a slice thereof) is requested, the API will fill a *Rcpp*-style **Vector** object with corresponding data values from the matrix. A request for a specific entry of the matrix will directly return the corresponding data value.

In the following text, we discuss some of the specifics of the *beachmat* API implementation. This includes the details of each matrix representation, its memory footprint and the computational time required for data access.

### Performance with simple matrices

By default, R stores matrices as one-dimensional arrays of length *N_r_N_c_*, where *N_r_* and *N_c_* are the number of rows and columns, respectively. This is done in column-major format, i.e., the matrix entry (*x, y*) corresponds to array element *x* + *N_r_y* (assuming zero-based indexing). We refer to this format as a “simple matrix”. The simple matrix is easy to manipulate and the time required for data access is linear with respect to the number of rows/columns (Supplementary Figure 1). However, its memory footprint is directly proportional to its length. For example, a double-precision matrix containing data for 10000 genes in each of one million cells would require 80 GB of RAM to store in memory. This is currently not possible for most workstations, instead requiring dedicated high-performance computing resources. Even smaller matrices will cause problems on systems with limited memory due to R’s copy-on-write semantics. Thus, the utility of simple matrices is limited to relatively small scRNA-seq data sets.

We compare the access speed of the *beachmat* API to that of a reference implementation using only *Rcpp*. Both row and column access via *beachmat* require 20-50% more time compared to the reference (Supplementary Figure 1). This is expected as *beachmat* is built on top of *Rcpp*, so the former cannot be faster than the latter. Another reason is that, at each row/column access, *beachmat* copies the matrix data into a **Vector** by default. The reference implementation avoids the overhead of creating a new copy by simply iterating across the original data. Our use of copying is deliberate as it ensures that the API is consistent across matrix representations – for example, file-based representations *must* copy the data to a new location in memory. Copying is also required for operations that involve transformations and/or re-ordering of data, as well as for libraries such as LAPACK that accept a pointer to a contiguous block of memory. Nonetheless, for read-only access to column data in a simple matrix, developers can direct *beachmat* to return an iterator directly to the start of the column. This avoids making a copy and allows for some optimization in certain scenarios.

### Performance with sparse matrices

The **dgCMatrix** class from the *Matrix* package stores sparse matrix data in compressed sparse column-orientated (CSC) format (Supplementary Figure 2). Consider that every non-zero entry in this matrix is characterized by a triplet: row *x*, column *y* and value *v*. To convert this into the CSC format, entries are sorted in order of increasing *x* + *N_r_y*. All entries with the same value of *y* are now grouped together in the ordered sequence. We refer to each column-based group as *G_y_*, the entries of which are sorted internally in order of increasing *x*. The representation is further compressed by discarding *y* from each triplet. All entries from the same column are at consecutive locations of the ordered sequence, so only the start position of *G_y_* on the sequence needs to be stored for each column. (The end position of one column is simply the start position of the next column.) This reduces memory usage to *s_I_N_c_* + (*s_I_* + *s_v_*)*N*_≠0_ where *s_I_* is the size of an integer, *s_v_* is the size of a single data element and *N*_≠0_ is the number of non-zero elements in the matrix. For double-precision matrices with many rows, sparse matrices will be more memory-efficient than their simpler “dense” counterparts if the density of non-zero elements is less than *≈* 66% (assuming 4-byte integers and 8-byte doubles).

The CSC format simplifies data access by imposing structure on the non-zero entries. When accessing a particular column *y*, all corresponding entries in *G_y_* can be quickly extracted by taking the relevant part of the ordered sequence. For low-density sparse matrices, column access via *beachmat* is even faster than access from simple matrices (Supplementary Figure 3). This is because only a few non-zero entries need to be copied – the rest of the **Vector** can be rapidly filled with zeroes. As the density of non-zero entries increases, column access becomes slower but is still comparable to that of simple matrices. We note that the *RcppArmadillo* package [5] also handles sparse matrices via the **SpMat** class. This provides faster column-level access than the *beachmat* API as no copying of data is performed – see above for a related discussion with simple matrices.

Row-level access is more difficult in the CSC format as entries in the same row do not follow a predictable pattern. If a row is requested, a binary search on *x* needs to be performed within *G_y_* for each column *y*, which requires an average time proportional to log(*N_r_*). In contrast, obtaining the next element in a row of a dense matrix can be done in constant time by jumping *N_r_* elements ahead on the one-dimensional array. To speed up row access for sparse matrices, we realized that the most common access pattern involves requests for consecutive rows. If row *r* is accessed, *beachmat* will loop over all columns and cache the index of the first value of *x* in *G_y_* that is not less than *r*. When *r* + 1 is accessed, we simply need to check if each of the indices should be incremented by one. This avoids the need to perform a new binary search and reduces the row access time substantially (Figure 2). Even when the row access pattern is random, we mitigate the time penalty by checking if the requested row is greater than or less than the previous row for which indices are stored. If greater, we use the stored indices to set the start of the binary search; if less, we use the indices to set the end of the search. This reduces the search space and the amount of computational work for large *N_r_*.

**Figure 2.**
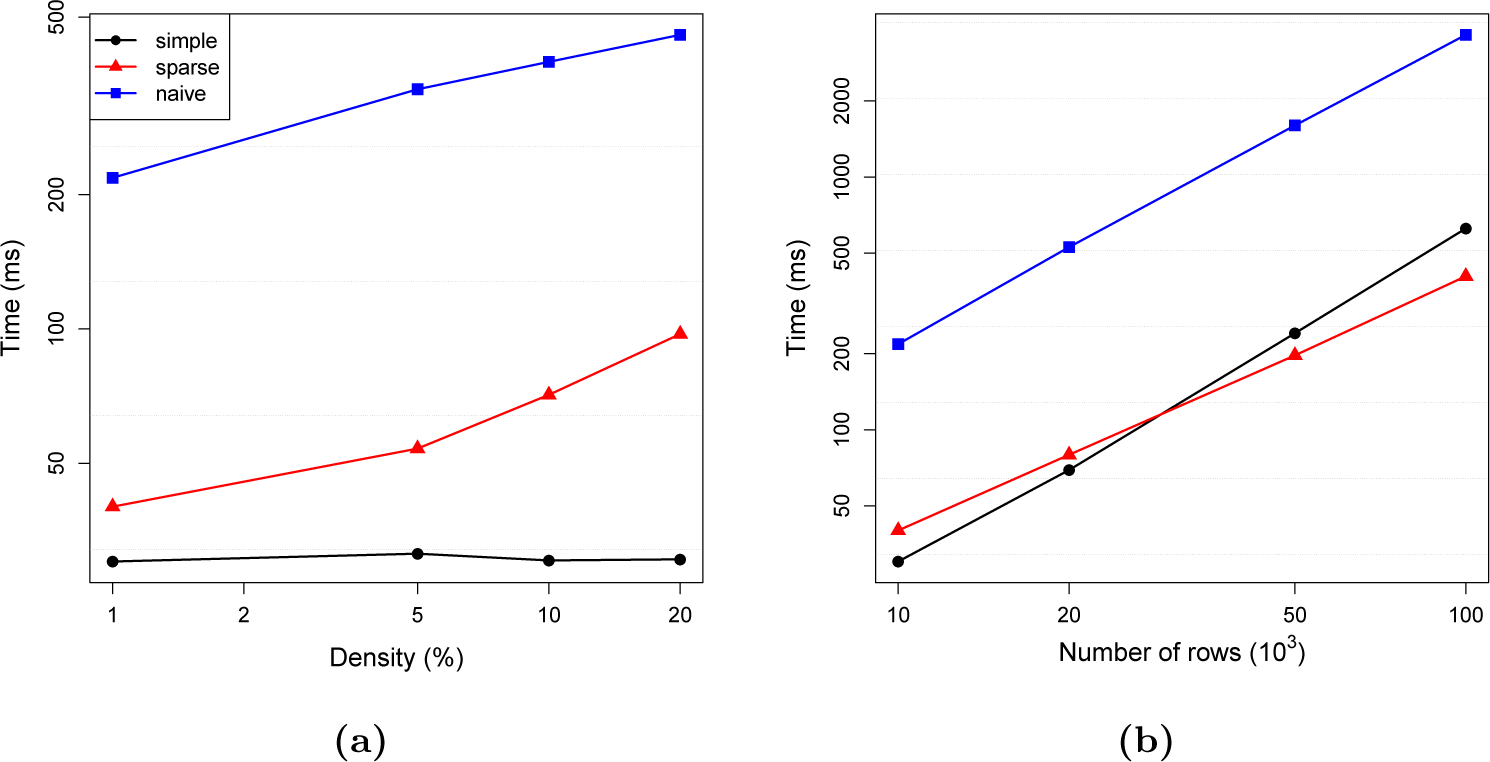
Row access times for CSC matrices using a naive binary search or the improved caching method in *beachmat*. For reference, access time for an equivalent simple matrix in *beachmat* (simple) is also shown. (a) Access times with respect to the density of non-zero entries, for a matrix with 10000 rows and 1000 columns. (b) Access times with respect to the number of rows, for a matrix with 1000 columns and 1% non-zero entries. Horizontal dotted lines represent 2-fold increases in time.

Despite these optimisations, row access with sparse matrices in *beachmat* remains slower than that with simple matrices (Figure 2a). This is not surprising as there is simply less work to do with dense matrices. The exception is with large matrices of low density (Figure 2b), where the cost of CPU cache misses due to large jumps exceeds the cost of handling both *x* and *v* per non-zero entry. Row access of either representation with *beachmat* is also faster than that of sparse matrices with *RcppArmadillo*. For example, for a 10000-by-1000 sparse matrix with 1% non-zero entries, *beachmat* with sparse matrices takes 39.8 milliseconds to access each row; *beachmat* with dense matrices takes 29.2 milliseconds; and *RcppArmadillo* takes 1921.4 milliseconds. These results motivate the use of *beachmat* for data access from sparse matrices.

For highly sparse data, it is also possible to design efficient algorithms that completely avoid processing the zeroes. To accommodate this, *beachmat* can be directed to return the indices and values of all non-zero elements in each row or column. In contrast, the default acccess methods will return all values of a row/column. This ensures that the API is consistent with non-sparse representations but is inefficient for operations where zeroes can be ignored. The user-level code can switch between different algorithms depending on whether the input matrix is sparse or dense. In general, we recommend this approach only for performance-critical parts of the code. Writing, testing and maintaining two different versions of the same code doubles the burden on the developer, which is precisely what *beachmat* was intended to avoid.

### Performance with HDF5-based matrices

For large, non-sparse matrices that do not fit into memory, the most obvious option is to store them on disk and load submatrices into memory as required. We consider the use of the hierarchical data format (HDF5) [23], which provides flexible and efficient storage of and access to large amounts of data in a filesystem-like format. Each large matrix is stored as a dataset in a HDF5 file on disk, while in memory, it is represented in R by a **HDF5Matrix** object from the *HDF5Array* package. The in-memory representation is very small – fewer than 3 kilobytes in size – and simply extracts data from disk upon request. Each **HDF5Matrix** instance provides methods to mimic a real matrix object and allows users to manipulate the matrix in real time without the need to load all of the data into memory. Compression of data in the HDF5 file also ensures that the on-disk footprint remains manageable throughout the course of the analysis.

The *beachmat* API supports row- and column-level access from a **HDF5Matrix** instance. Specifically, *beachmat* directly accesses data from the underlying HDF5 file through the official HDF5 C++ API, which has been stored in the *Rhdf5lib* package for portability. This means that even very large data sets can be accessed in C++, using the same code for simple and sparse matrix representations. However, data access is inevitably slower than that from a simple matrix as the data need to be read from disk at regular intervals. We observed a 20-fold increase in the time required to access each column and a 40-fold increase in the time required to access each row (Figure 3). This suggests that the **HDF5Matrix** representation should be used sparingly – if possible, smaller data sets should use alternative in-memory representations for faster access.

**Figure 3.**
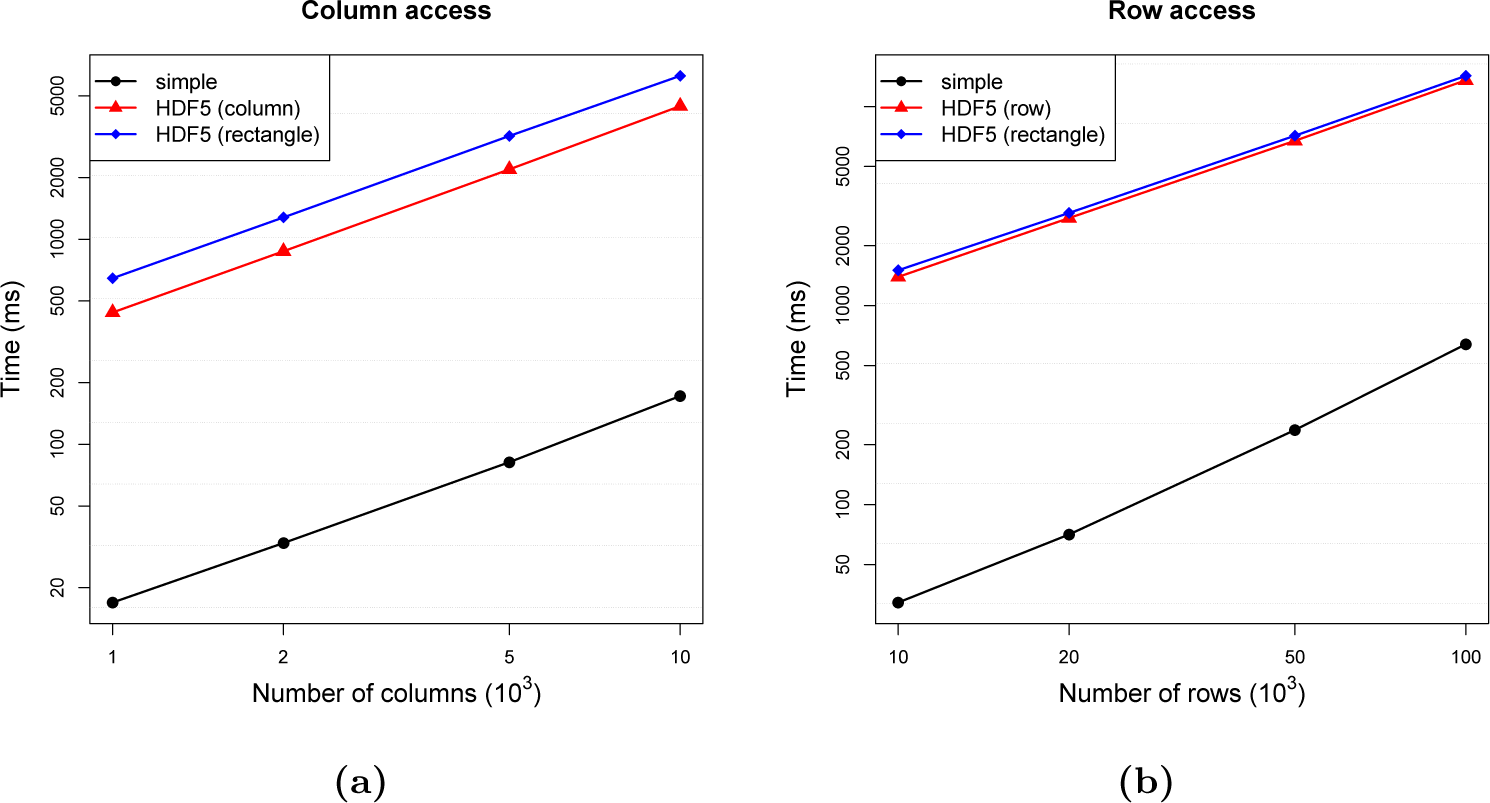
Access times for a HDF5-backed matrix using column/row-chunking or rectangular 100 *×* 100 chunks. Times for a simple matrix are shown for comparison. (a) Column access time with respect to the number of columns, for a dense matrix with 10000 rows. (b) Row access time with respect to the number of rows, for a dense matrix with 1000 columns. Horizontal dotted lines represent 2-fold increases in time.

A key determinant of the performance of the **HDF5Matrix** representation is the layout of data in the HDF5 file. There are two layout choices for large matrices: contiguous or chunked. In the contiguous layout, raw data are flattened into a one-dimensional array analogous to column-major storage of simple matrices in memory. In the chunked layout, data are arranged into “chunks” of a pre-defined size. For example, in a row-chunked layout, each chunk would correspond to a row of the matrix. Each chunk is always read (or written) in its entirety, even when only a portion of the chunk is requested. Chunking is required for fast access to data, *provided that the layout is consistent with the expected access pattern*. For example, a row-chunked layout allows fast access to each row, as only one disk read is required to obtain the chunk for each row. However, access to any given column is very slow, as the value of each element in the column must be obtained by performing a disk read for every row chunk in its entirety, i.e., *N_r_* reads in total. In practice, both row and column accesses are often required (e.g., to access gene- and cell-level scRNA-seq data), which means that the file layout must be carefully chosen to allow for these orthogonal access patterns.

The choice of file layout is the responsibility of the process that constructs the HDF5 file. This can be the original data provider; the developer whose function returns a **HDF5Matrix**; or a user who coerces their data into a **HDF5Matrix**. As such, the chunking scheme is generally outside of *beachmat*’s control, preventing the API from automatically choosing the optimal layout for the requested access pattern. Nonetheless, for a given layout, *beachmat* will dynamically resize the HDF5 chunk cache to speed up access to consecutive rows or columns – see Additional Note 1 for more information. This permits fast extraction of both row- and column-based data from the same layout (Figure 3, Supplementary Figures 4-5, Additional Note 2). *beachmat* also provides a function to convert an existing HDF5 file to pure row- or column-based chunks (Additional Note 2), which performs optimally for random row and column access, respectively.

Another benefit of chunking is that the data in each chunk can be compressed using filters such as ZLIB and SZIP. This decreases the size of the HDF5 file by at least 4-fold for dense matrices (80 MB to 19 MB for a 10000-by-1000 double-precision matrix), with even greater gains for sparse data (9 MB for the same matrix with 1% non-zero entries). The use of smaller files reduces the risk that disk space will be exceeded during the course of an analysis. This is important when many on-disk matrices need to be constructed, e.g., to store transformed expression values or intermediate results.

### Other matrix types

While the matrix representations described above are the most commonly used for storing scRNA-seq data, *beachmat* can be easily extended to other representations. For example, the packed symmetric representation from the *Matrix* package only stores the upper or lower half of a symmetric matrix. This provides an efficient representation of distance matrices, which are often used to cluster cells based on their expression profiles. *beachmat* supports row and column access of data from packed symmetric matrices through the same interface that is used for the other representations.

*beachmat* also supports data access from **RleMatrix** instances from the *DelayedArray* package. The **RleMatrix** stores its values as a column-major run-length encoding, where stretches of the same value in the one-dimensional array are stored as a single run. This reduces memory usage in a more general manner than a sparse matrix, especially for matrices with many small but non-zero counts. As with CSC matrices, *beachmat* caches the row indices to speed up consecutive row access.

Another option for storing matrices on disk is to use the *bigmemory* package [10]. This constructs an in-memory **big.matrix** object that contains external pointers pointing to an on-disk representation. In *beachmat*, we have deliberately chosen **HDF5Matrix** rather than **big.matrix** due to the standardized nature of the HDF5 specification and portability of HDF5 files across systems. Nonetheless, we note that it is simple to extend *beachmat* to accept **big.matrix** inputs if required.

### Storing matrix output

In addition to accessing data in existing matrices, the *beachmat* API allows values generated in C++ to be stored in various matrix representations for output to R. For integer, logical, double-precision and character data, simple and HDF5-backed matrices can be constructed that are indistinguishable from those generated in R. Logical and double-precision data can also be stored in CSC format, where only true or non-zero values are retained in **lgCMatrix** or **dgCMatrix** instances, respectively. (The *Matrix* package does not support sparse integer or character matrices, so these are ignored.)

The output representation can either be explicitly specified in the code, or it can be automatically chosen to match some input representation. To illustrate, consider a C++ function that accepts a matrix as input and returns a matrix of similar dimensions. If the input is in the simple matrix format, one might assume that there is enough memory to also store the output in the simple format; whereas if the input is a **HDF5Matrix**, one could presume that the output would be similarly large, thus requiring a **HDF5Matrix** representation for the results. This means that results of processing in C++ can be readily returned in the most suitable representation for manipulation in R.

Note that the layout considerations described for read access to HDF5-backed input are equally applicable to HDF5-backed output. If rows/columns are to be filled consecutively, we suggest using rectangular chunks that are proportional to the dimensions of the matrix (Additional Note 1). For random write access, pure column-or row-based chunks are more suitable. Chunk dimensions can be specified directly with the *beachmat* API; or by using functions from the *HDF5Array* package to set the global chunking dimensions in R, which will be respected by *beachmat* in the C++ code.

## Performance of *beachmat* on real data sets

### Access times for a small brain data set

We evaluated the performance of *beachmat* with the different matrix representations on real data, using the count matrix from a scRNA-seq study of the mouse brain [24]. This data set contains integer expression values for 19972 genes in each of 3005 cells, of which 18% are non-zero. We note that this is not a particularly large matrix, especially in the context of droplet-based experiments that routinely generate data for tens of thousands of cells. However, its size ensures that each of the matrix representations – including those that are stored in memory – can be easily evaluated and compared.

The performances of the different representations on the brain data set largely recapitulate the results with simulated data. Row and column accesses from a simple matrix are the fastest, followed by accesses from a sparse matrix (Figure 4). HDF5-backed matrices provide the slowest access but also the smallest memory footprint (2 KB, compared to 480 MB for simple matrices and 130 MB for sparse matrices). The on-disk size of the HDF5 file is also relatively small, requiring only 16-20 MB of space for each **HDF5Matrix** instance. These results demonstrate that the strengths and weaknesses of the different representations are recapitulated with real data.

**Figure 4.**
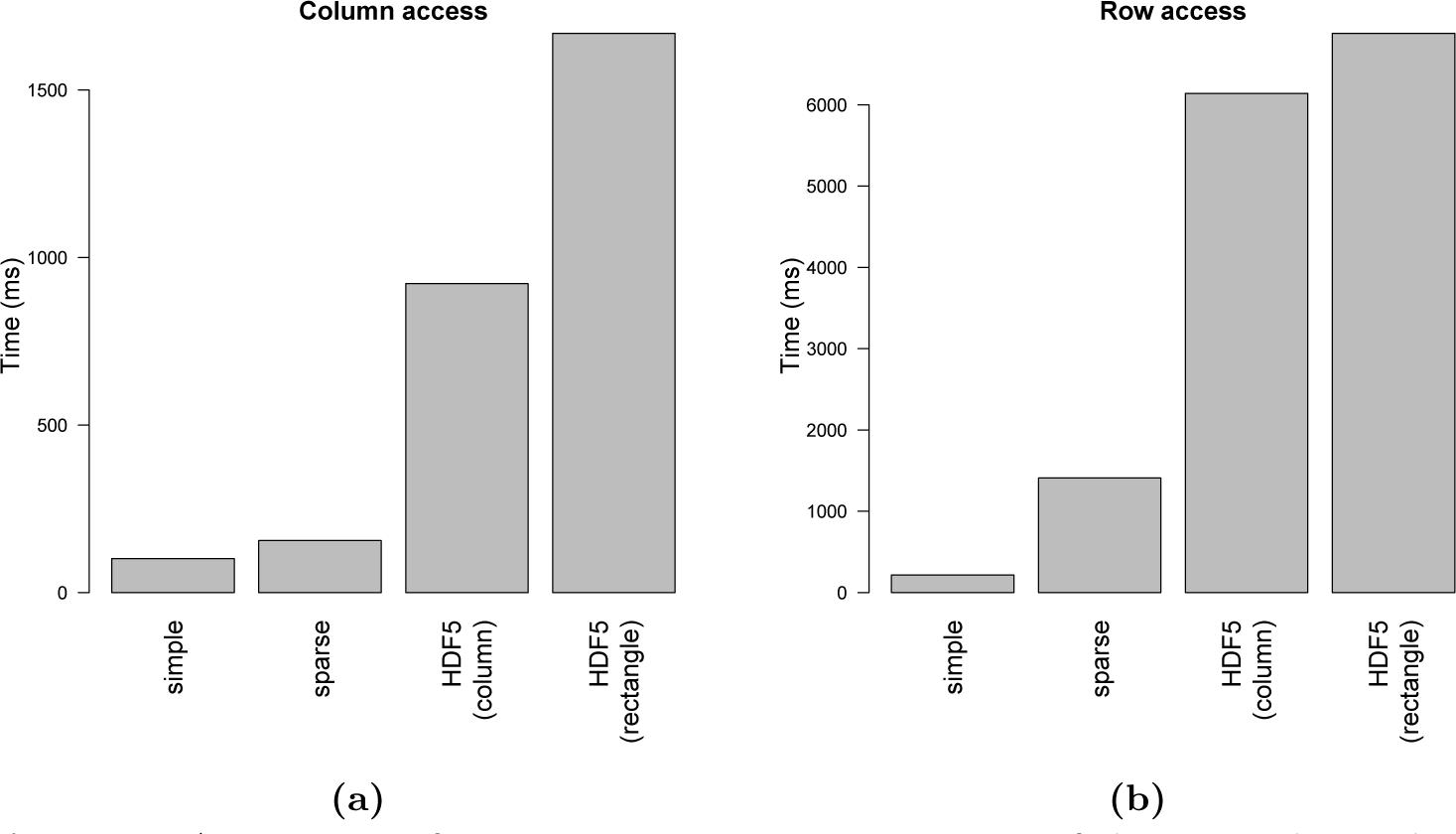
Access times for various matrix representations of the mouse brain data set from Zeisel *et al.* [24]. (a) Column access time for each representation, based on calculation of column sums. For **HDF5Matrix**, a column-chunked layout and a rectangular 200 *×* 200 layout were tested. (b) Row access time for each representation. For **HDF5Matrix**, a row-chunked layout and a rectangular 200 *×* 200 layout were tested. Heights represent the average of 10 repeated timings; standard errors were negligible and not shown.

#### Analysis of the very large 10X data set

To demonstrate the utility of *beachmat* for faciliting analyses of large data sets, we converted several functions in the *scater* [17] and *scran* packges [14] to use the *beachmat* API in their C++ code. We applied these functions to the 1 million neuron data set from 10X Genomics (see Methods). First, we called cell cycle phase with the cyclone method [21]. The vast majority of cells were identified as being in G1 phase (Figure 5a), consistent with the presence of differentiated neurons that are not actively cycling. Next, we applied the deconvolution method [13] to compute size factors to normalize for cell-specific biases. The size factor generally correlated well with the library size for each cell (Figure 5b). Deviations were observed for a small number of cells, consistent with composition effects [20] caused by differential expression between cell subpopulations.

**Figure 5.**
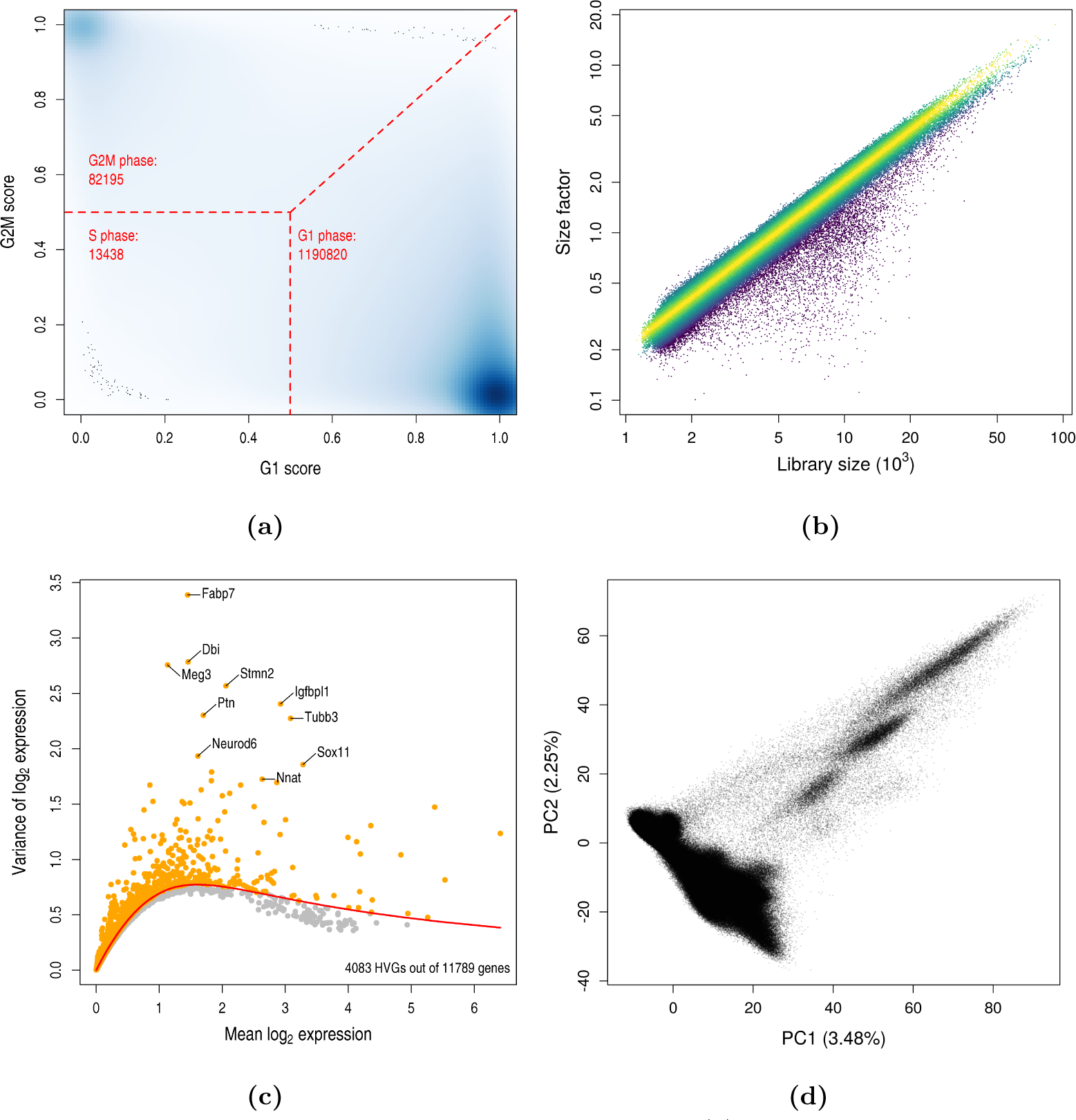
Analysis of the 10X million neuron data set. (a) Cell cycle phase assignment, based on the G1 and G2M scores reported by **cyclone**. The intensity of colour is proportional to the density of cells at each plot location. Dashed lines indicate the score boundaries corresponding to each phase, and the number of assigned cells is also shown for each phase. (b) Size factor for each cell from the deconvolution method, plotted against the library size. Cells were coloured according to the deviation from the median log-ratio of the size factor to the library size for all cells. (c) Variance of the normalized log-expression values for each gene, plotted against the mean log-expression. The red line indicates the mean-dependent trend fitted to all genes. Orange points correspond to highly variable genes detected at a false discovery rate of 5%, with the top 10 genes highlighted. (d) PCA plot generated from the HVG expression profiles of all cells. The variance explained by each of the first two principal components is shown in brackets.

We then detected highly variable genes (HVGs) based on the variance of the log-normalized expression values [14]. This was performed while blocking on the sequencing library of origin for each cell, to regress out technical factors of variation unrelated to biological heterogeneity. We identified a number of HVGs (Figure 5c), including genes involved in neuronal differentiation and function such as *Neurod6* [11] and *Sox11* [2]. Finally, we performed dimensionality reduction on the HVG expression profiles for all cells using principal components analysis (PCA). This showed clear substructure in the first two principal components (PCs), reflecting the diversity of cell types in the mouse brain. Indeed, once PCA has been performed, the first 10-20 PCs for each cell can be used as a summary of its expression profile. This can be stored in memory as a simple matrix and supplied directly to other R functions for further processing, provided the underlying algorithms are scalable with the number of cells.

While a full characterisation of this data set is outside the scope of this paper, it is clear that we can proceed through many parts of the scRNA-seq analysis pipeline using *beachmat*-driven C++ functions. By taking advantage of the file-backed **HDF5Matrix**, this analysis can be conducted in reasonable time on a desktop with modest specifications (see Methods). In particular, the incoporation of the *beachmat* API only required modest modifications to the existing C++ code for scRNA-seq data analysis. Obtaining this level of functionality without *beachmat* would be much more difficult.

### Discussion

The popularity of the R programming language stems, in part, from the ease of its extensibility. Packages can be easily developed by the research community to implement cutting-edge algorithms for new sources of data. The increasing number of packages designed to analyze scRNA-seq data (13 on Bioconductor at time of writing) provides a case in point. Here, we describe the *beachmat* package, which provides a common API for data access from a variety of R matrix representations in C++. This simplifies package development and improves interoperability within the R/Bioconductor ecosystem, by enabling arbitrary C++ code to accept many different matrix inputs without any further effort on the part of the developer. While we have focused on scRNA-seq in this paper, analyses of other large matrices (e.g., genome-wide contact matrices in Hi-C data [15]) may also benefit from *beachmat*-driven code.

As we have shown, each matrix representation has specific strengths and weaknesses for data access. Matrices that occupy more memory generally provide faster access, as data do not need to be unpacked or retrieved from disk. Obviously, though, this may not be practical for large data sets. Sparse matrix representations are not effective if sparsity-destroying operations (e.g., mean-centering during PCA) are applied. Even high-performance computing resources have their limits, especially in academic environments with many users where high-resource jobs are difficult to schedule. In such cases, it may be preferable to sacrifice speed for reduced memory consumption by using a file-backed representation such as the **HDF5Matrix** class. By incorporating *beachmat* into the C++ code, an R package can dynamically accept different matrix types appropriate for the size of the data set and computing environment.

An alternative to using *beachmat* is to write C++ code for one matrix representation (usually simple matrices) and apply it to chunks of a given input matrix. Each chunk is coerced into the chosen representation and the C++ code is applied to the coerced object. After looping across all chunks, the chunk-wise results are combined to obtain the final result for the entire matrix. However, this hybrid approach requires extensive coordination between R and C++ to keep track of the chunk that is being processed, to monitor intermediate variables that persist between chunks, and to combine the results in an appropriate manner. The need to ensure that R and C++ are interacting correctly at multiple points imposes a significant burden on the developer. Computational efficiency is also reduced by the use of R loops, multiple matrix coercions and repeated C++ function calls. The *beachmat* API provides a natural solution by moving the entire procedure into C++, simplifying development and maintenance.

It is straightforward to integrate *beachmat* into C++ code in existing R packages. Our modifications to *scran* and *scater* have enabled the analysis of a very large scRNA-seq data set in low-memory environments using file-backed representations, without compromising speed for smaller data sets that can be held fully in memory. We anticipate that *beachmat* will be useful to developers of computationally intensive bioinformatics methods that need to access data from different matrices. Given the infrastructure that is now available for handling large data sets, it is fair to say that the rumours of R’s demise (Supplementary Figure 6) have been greatly exaggerated.

## Code availability

*beachmat* is available as part of version 3.6 of the Bioconductor project (https://bioconductor.org/packages/beachmat). The *scran* and *scater* packages that were modified to support *beachmat* can also be installed from Bioconductor. All code used to perform the simulations and real data analyses are available on Github (https://github.com/LTLA/MatrixEval2017).

## Author contributions

ATLL conceived and implemented the *beachmat* C++ API, performed the data access simulations and analyzed the 10X data set. HP adapted the *HDF5Array* and *DelayedArray* packages to interface with *beachmat*. MLS ported the HDF5 C++ API into *Rhdf5lib*, to support HDF5 read/write access in *beachmat*.

## Acknowledgements

We thank Martin Morgan, Andrew McDavid, Peter Hickey, Raphael Gottardo, Mike Jiang, Vince Carey, John Readey and other members of the Bioconductor single-cell big data working group for useful discussions. We thank Davis McCarthy for his assistance with incorporating *beachmat* into *scater*. We also thank John Marioni for helpful comments on the manuscript.

## Methods

### Access timings with simulated data

Double-precision dense matrices of a specified dimension were filled with values sampled from a standard normal distribution. By default, all dense matrices were constructed in the simple format. Double-precision CSC matrices of a specified dimension and density were generated as **dgCMatrix** instances, using the **rsparsematrix** function from the *Matrix* package. Conversion of each matrix object into other representations, if necessary, was performed prior to timing access with the C++ APIs.

To time row and column access with *beachmat* and other APIs, we wrote a C++ function that computes the sum of values in each row or column, respectively. Calculation of the row/column sums ensures that each entry of the row/column is visited in order to use its value. This means that each API has to do some work, avoiding trivially fast approaches where a pointer or iterator is returned to the start of the column/row. Summation is also simple enough that the access time of the API will still constitute a major part of the overall time spent by the function

Timings were performed in R using the **system.time** function on a call to each C++ function via .**Call**. This was repeated 10 times with new matrices, and the average time and standard error were computed for each access method. Row and column access times were evaluated with respect to the numbers of rows and columns, respectively. For sparse matrices, times were also recorded with respect to the density of non-zero entries. In all cases, standard errors were negligible and not plotted for clarity.

### Access timings with the brain data

scRNA-seq data from the mouse brain study [24] were obtained as a count matrix from http://linnarssonlab.org/cortex/. Counts were read into R and converted into a double-precision simple matrix, a **dgCMatrix** or a **HDF5Matrix**. For each representation, timings of the calculation of row or column sums were performed as previously described. This was repeated 10 times to obtain an average time. Column- and row-wise chunking were used for timing column and row access, respectively, of a **HDF5Matrix**.

### Analysis of the 1 million neuron data set

We downloaded the 1 million neuron data set from the 10X Genomics website (https://support.10xgenomics.com/single-cell-gene-expression/datasets/1.3.0/1M_neurons, obtained 15 June 2017). We used the *TENxGenomics* package (https://github.com/mtmorgan/TENxGenomics) to compute metrics for each cell, including the total number of unique molecular identifiers (UMIs) and the total number of expressed genes (i.e., with at least one UMI). Low-quality cells were identified as those with log-total UMI counts or log-numbers of expressed genes that were more than three median absolute deviations below the median value for all cells in the same sequencing library. After filtering out the low-quality cells, we converted the count data into a **HDF5Matrix** with column-based chunks using the *HDF5Array* package.

To call cell cycle phase, we applied the cyclone method [21] from the *scran* package using the pre-defined mouse classifier. This was performed over three cores to reduce computational time. To compute size factors, we used the **computeSumFactors** method after splitting the data set into chunks of 2000-3000 cells. Cells in each chunk were normalized using the deconvolution method [13], and size factors were calibrated across chunks by normalizing the chunk-specific pseudo-cells. The size factor for each cell was used to compute normalized log-expression values [14], which were represented in a new **HDF5Matrix** object. Note that, when computing the size factors, we only used genes with an average count above 0.1 (as calculated by the **calcAverage** function in *scater*). This ensures that the pooled expression profiles will not be dominated by zeros.

To identify highly variable genes, we used the **trendVar** and **decomposeVar** functions from *scran*. We computed the variance of the normalized log-expression values for each gene while blocking on the sequencing library of origin. We fitted a mean-dependent trend to the variances to model the mean-variance relationship. Assuming that most genes were not highly variable, we tested whether the residual from the trend for each gene was significantly greater than zero. Highly variable genes were identified as those with a significantly non-zero component at a FDR of 5%. To reduce computational time, the severity of the multiple testing correction and to avoid discreteness when fitting the trend, we only considered genes with an average count above 0.01.

We used a simple approach to perform PCA on a very large matrix. After subsetting the expression matrix by the detected HVGs, we mean-centred and standardized the expression vector for each gene. We randomly selected 10000 cells and coerced the expression profiles into a simple matrix. This was used as input into the **prcomp** function to obtain the loading vectors. We then projected all cells onto the space defined by the first two loading vectors to obtain PC1 and PC2 coordinates for all cells. This approach assumes the selected 10000 cells provide a good representation of the variance structure in the full population, allowing the approximate loading vectors to be obtained by PCA on the smaller matrix. More sophisticated strategies such as randomized PCA [8] could also be used, but a correct and efficient implementation of such algorithms for **HDF5Matrix** objects is beyond the scope of this paper.

### Details of the computing environment

All timings and analyses were performed on a Dell OptiPlex 790 desktop with an Intel Core i5 processor and 8 GB of RAM, running Ubuntu 14.04.5 with R 3.4.0 and Biocondcutor 3.6. On this machine, the 10X data analysis required approximately 4 hours for quality control and construction of the **HDF5Matrix**; 22 hours for cell cycle phase assignment; 8 hours for calculation of size factors and generation of normalized expression values; 10 hours for detecting highly variable genes; and 8 hours for PCA. Only the phase assignment step was run on multiple cores.

## References

1. D. Bates and M. Maechler. Matrix: Sparse and Dense Matrix Classes and Methods, 2017. R package version 1.2-10.

2. M. Bergsland, M. Werme, M. Malewicz, T. Perlmann, and J. Muhr. The establishment of neuronal properties is controlled by Sox4 and Sox11. Genes Dev., 20(24):3475–3486, Dec 2006.

3. J. D. Buenrostro, B. Wu, U. M. Litzenburger, D. Ruff, M. L. Gonzales, M. P. Snyder, H. Y. Chang, and W. J. Greenleaf. Single-cell chromatin accessibility reveals principles of regulatory variation. Nature, 523(7561):486–490, Jul 2015.

4. D. Eddelbuettel and R. François. Rcpp: Seamless R and C++ integration. Journal of Statistical Software, 40(8):1–18, 2011.

5. D. Eddelbuettel and C. Sanderson. RcppArmadillo: Accelerating R with high-performance C++ linear algebra. Computational Statistics and Data Analysis, 71:1054–1063, March 2014.

6. R. C. Gentleman, V. J. Carey, D. M. Bates, B. Bolstad, M. Dettling, S. Dudoit, B. Ellis, L. Gautier, Y. Ge, J. Gentry, K. Hornik, T. Hothorn, W. Huber, S. Iacus, R. Irizarry, F. Leisch, C. Li, M. Maechler, A. J. Rossini, G. Sawitzki, C. Smith, G. Smyth, L. Tierney, J. Y. Yang, and J. Zhang. Bioconductor: open software development for computational biology and bioinformatics. Genome Biol., 5(10):R80, 2004.

7. D. Grun and A. van Oudenaarden. Design and analysis of single-cell sequencing experiments. Cell, 163(4):799–810, Nov 2015.

8. N. Halko, P. G. Martinsson, and J. A. Tropp. Finding structure with randomness: Probabilistic algorithms for constructing approximate matrix decompositions. SIAM Review, 53(2):217–288, 2011.

9. W. Huber, V. J. Carey, R. Gentleman, S. Anders, M. Carlson, B. S. Carvalho, H. C. Bravo, S. Davis, L. Gatto, T. Girke, R. Gottardo, F. Hahne, K. D. Hansen, R. A. Irizarry, M. Lawrence, M. I. Love, J. MacDonald, V. Obenchain, A. K. Oleś, H. Pages, A. Reyes, P. Shannon, G. K. Smyth, D. Tenenbaum, L. Waldron, and M. Morgan. Orchestrating high-throughput genomic analysis with Bioconductor. Nat. Methods, 12(2):115–121, Feb 2015.

10. M. J. Kane, J. Emerson, and S. Weston. Scalable strategies for computing with massive data. Journal of Statistical Software, 55(14):1–19, 2013.

11. J. N. Kay, P. E. Voinescu, M. W. Chu, and J. R. Sanes. Neurod6 expression defines new retinal amacrine cell subtypes and regulates their fate. Nat. Neurosci., 14(8):965–972, Jul 2011.

12. A. M. Klein, L. Mazutis, I. Akartuna, N. Tallapragada, A. Veres, V. Li, L. Peshkin, D. A. Weitz, and M. W. Kirschner. Droplet barcoding for single-cell transcriptomics applied to embryonic stem cells. Cell, 161(5):1187–1201, May 2015.

13. A. T. Lun, K. Bach, and J. C. Marioni. Pooling across cells to normalize single-cell RNA sequencing data with many zero counts. Genome Biol., 17:75, Apr 2016.

14. A. T. Lun, D. J. McCarthy, and J. C. Marioni. A step-by-step workflow for low-level analysis of single-cell RNA-seq data with Bioconductor. F1000Res, 5:2122, 2016.

15. A. T. Lun, M. Perry, and E. Ing-Simmons. Infrastructure for genomic interactions: Bioconductor classes for Hi-C, ChIA-PET and related experiments. F1000Res, 5:950, 2016.

16. E. Z. Macosko, A. Basu, R. Satija, J. Nemesh, K. Shekhar, M. Goldman, I. Tirosh, A. R. Bialas, N. Kamitaki, E. M. Martersteck, J. J. Trombetta, D. A. Weitz, J. R. Sanes, A. K. Shalek, A. Regev, and S. A. McCarroll. Highly parallel genome-wide expression profiling of individual cells using nanoliter droplets. Cell, 161(5):1202–1214, May 2015.

17. D. J. McCarthy, K. R. Campbell, A. T. Lun, and Q. F. Wills. Scater: pre-processing, quality control, normalization and visualization of single-cell RNA-seq data in R. Bioinformatics, 33(8):1179–1186, Apr 2017.

18. R Core Team. R: A Language and Environment for Statistical Computing. R Foundation for Statistical Computing, Vienna, Austria, 2017.

19. A. Regev, S. Teichmann, E. S. Lander, I. Amit, C. Benoist, E. Birney, B. Bodenmiller, P. Campbell, P. Carninci, M. Clatworthy, H. Clevers, B. Deplancke, I. Dunham, J. Eberwine, R. Eils, W. Enard, A. Farmer, L. Fugger, B. Gottgens, N. Hacohen, M. Haniffa, M. Hemberg, S. K. Kim, P. Klenerman, A. Kriegstein, E. Lein, S. Linnarsson, J. Lundeberg, P. Majumder, J. Marioni, M. Merad, M. Mhlanga, M. Nawijn, M. Netea, G. Nolan, D. Pe’er, A. Philipakis, C. P. Ponting, S. R. Quake, W. Reik, O. Rozenblatt-Rosen, J. R. Sanes, R. Satija, T. Shumacher, A. K. Shalek, E. Shapiro, P. Sharma, J. Shin, O. Stegle, M. Stratton, M. J. T. Stubbington, A. van Oudenaarden, A. Wagner, F. M. Watt, J. S. Weissman, B. Wold, R. J. Xavier, and N. Yosef. The human cell atlas. bioRxiv, 2017.

20. M. D. Robinson and A. Oshlack. A scaling normalization method for differential expression analysis of RNA-seq data. Genome Biol., 11(3):R25, 2010.

21. A. Scialdone, K. N. Natarajan, L. R. Saraiva, V. Proserpio, S. A. Teichmann, O. Stegle, J. C. Marioni, and F. Buettner. Computational assignment of cell-cycle stage from single-cell transcriptome data. Methods, 85:54–61, Sep 2015.

22. S. A. Smallwood, H. J. Lee, C. Angermueller, F. Krueger, H. Saadeh, J. Peat, S. R. Andrews, O. Stegle, W. Reik, and G. Kelsey. Single-cell genome-wide bisulfite sequencing for assessing epigenetic heterogeneity. Nat. Methods, 11(8):817–820, Aug 2014.

23. The HDF Group. Hierarchical Data Format, version 5, 1997-2017. http://www.hdfgroup.org/HDF5/.

24. A. Zeisel, A. B. Munoz-Manchado, S. Codeluppi, P. Lonnerberg, G. La Manno, A. Jureus, S. Marques, H. Munguba, L. He, C. Betsholtz, C. Rolny, G. Castelo-Branco, J. Hjerling-Leffler, and S. Linnarsson. Brain structure. Cell types in the mouse cortex and hippocampus revealed by single-cell RNA-seq. Science, 347(6226):1138–1142, Mar 2015.

25. G. X. Zheng, J. M. Terry, P. Belgrader, P. Ryvkin, Z. W. Bent, R. Wilson, S. B. Ziraldo, T. D. Wheeler, G. P. McDermott, J. Zhu, M. T. Gregory, J. Shuga, L. Montesclaros, J. G. Underwood, D. A. Masquelier, S. Y. Nishimura, M. Schnall-Levin, P. W. Wyatt, C. M. Hindson, R. Bharadwaj, A. Wong, K. D. Ness, L. W. Beppu, H. J. Deeg, C. McFarland, K. R. Loeb, W. J. Valente, N. G. Ericson, E. A. Stevens, J. P. Radich, T. S. Mikkelsen, B. J. Hindson, and J. H. Bielas. Massively parallel digital transcriptional profiling of single cells. Nat Commun, 8:14049, Jan 2017.

